# Contrasting impacts of environmental variability on the breeding biology of two sympatric small procellariiform seabirds in south-eastern Australia

**DOI:** 10.1101/2021.04.19.440428

**Authors:** Yonina H. Eizenberg, Aymeric Fromant, Arnaud Lec’hvien, John P.Y. Arnould

## Abstract

Seabirds play a vital role in marine ecosystems and are determinant sentinels of the productivity of their environments. The long-term study of their breeding biology and their responses to environmental variations can be used to monitor the effects of climate change on marine fauna. However, the ecological and physiological differences among seabirds induce a large range of responses complicating our understanding of the effects of environmental changes on marine ecosystems. The present study investigated the impact of environmental variability on breeding biology in two sympatric small Procellariiform species, the fairy prion (*Pachyptila turtur*) and the common diving petrel (*Pelecanoides urinatrix*), over four reproductive seasons (2017-2020) in Bass Strait, south-eastern Australia. Marine heatwaves had a negative effect on chick growth, breeding success, and induced a delay in laying dates in both species. While fairy prions maintained a relatively high breeding success and broadly constant breeding phenology, common diving petrels delayed the start of the breeding season by up to 50 days and experienced dramatic collapses in breeding success in years of high marine heat wave occurrence. The high wing loading and absence of stomach oils in the common diving petrel are likely to have limited the capacity of this species to increase foraging effort in years of low food availability.

## Introduction

Seabirds are top-order predators with a global annual consumption of 69.8 million tonnes of marine biomass [1] comprised mainly of fish, crustaceans and cephalopods [2]. Correspondingly, seabirds play a vital role in the trophodynamics of marine food-webs and are essential for maintaining ecosystem function [3]. Consequently, seabirds are recognised as marine indicator species [1, 4] and knowledge of their breeding biology and reproductive success can provide valuable information on ocean health. This information can be used to identify changes in oceanic conditions and highlight potential threats that seabirds face [5].

Marine ecosystems are characteristically highly spatially and temporally variable [6, 7] and seabirds have evolved to be acutely adapted in their foraging strategy to maximise prey consumption for successful breeding [8]. However, throughout the world, seabirds are currently facing substantial immediate threats. Due to such factors as direct and indirect competition from fisheries, bycatch deaths in commercial fisheries, oil and plastics pollution and loss of breeding habitat [9–12], 28% of the 346 seabird species are currently listed as threatened [1, 13]. In addition, through changes to marine productivity, significant alterations to ocean temperatures [13] and currents [14] leading to shifts in prey species distribution, abundance and availability [15, 16], the effects of global climate change are considered the most pervasive and wide-ranging of impacts, affecting 40% of threatened seabirds [1, 5].

During the breeding season, seabirds adopt a central place foraging strategy and, therefore, are restricted in their ability to range long distances in search of prey resources [17, 18]. Consequently, they are susceptible to factors impacting the local distribution of their prey [13, 19]. With rapid changes associated with climate shifts, species may not be able to adjust their timing of breeding and foraging behaviour, leading to a mismatch of prey abundance to the energy requirements of chick-rearing [20–22]. As a consequence, modifications in behaviour and range shifts may occur when a species can no longer tolerate current food or habitat conditions [23–25].

Understanding how species may adapt to such changes can be obtained by observing their responses to environmental variability at the extent of their ranges [26–28]. More vulnerable to population decline than other seabird orders [28], Procellariiformes generally display a rigid breeding phenology (Chambers et al. 2014), experience late sexual maturity, lay a single egg each season and have slow chick growth [29–31]. In particular, small Procellariiformes (< 140 g) have a more limited foraging range and are, therefore, expected to be the most affected by environmental fluctuations. Although diverse, wide-ranging, contributing to a large biomass, and threatened by environmental changes, little is known about small Procellariiformes and their adaptation capacity [32].

The fairy prion (FP, *Pachyptila turtur*) and common diving petrel (CDP, *Pelecanoides urinatrix*) are two species of small Procellariiformes ubiquitous in temperate and subantarctic regions of the Southern Hemisphere [33]. Although both are similar sized (110 - 140 g), these species differ in biology and foraging strategy. Adult FP produce and feed stomach oil to chicks which enables fasting for long periods, allowing adults to have a greater foraging range during a time of increased prey dispersal [34]. Adult CDP do not produce stomach oil and, consequently, are required to feed chicks more regularly, limiting their foraging range during breeding [34, 35]. In addition, FP feed on the ocean surface while CDP may forage at depths of up to 30 m [36], potentially facilitating avoidance of interspecific competition for food [37].

Both species breed sympatrically at the northern extent of their range within Bass Strait, south-eastern Australia [38]. Bass Strait is recognised as a region of low marine productivity [39] and is expected to experience significant warming and changes to oceanic currents in the coming decades (IPCC, 2019) with potentially major impacts on the diversity, abundance and availability of prey resources [40, 41]. Considering 60% of Australian seabirds breed in Bass Strait [42], FP and CDP are especially at risk of interspecific competition for food in a changing ecosystem [33]. Relatively little is known of the ecology of these predominantly subantarctic species in south-eastern Australia [43, 44]. Such information is crucial for understanding how they may respond to the rapidly changing conditions in the area. Furthermore, knowledge of how current environmental variability impacts these species at the northern extent of their range is necessary for predicting how these, and other, small Procellariiformes will respond to the anticipated extreme environmental changes throughout subantarctic regions. The aims of the present study, therefore, were to: 1) determine breeding phenology; 2) assess chick growth and indices of breeding success; and 3) examine environmental factors influencing these parameters in FP and CDP nesting in northern Bass Strait.

## Materials and Methods

This study was conducted in accordance with the regulations of Deakin University Animal Ethics Committee (Approval B16-2017) and in accordance with the Department of Environment, Water, Land and Planning (Victoria) Wildlife Research Permit 10008452. Fieldwork was conducted over four consecutive breeding seasons (2017/18-2020/21) at Kanowna Island (39°09’S, 146°18’E) in northern Bass Strait, south-eastern Australia (Fig.1). Kanowna Island, a granite outcrop vegetated with coastal tussock grass (*Poa poiformis*), hosts *ca2000* FP and *ca500* CDP breeding pairs [43]. Since both species breed in subterranean burrows, the nest chamber was accessed by a short artificial tunnel and covered with a removable stone lid. This access system reduced the disturbance of the natural tunnel and facilitated rapid access to the birds [45].

**Figure 1:**
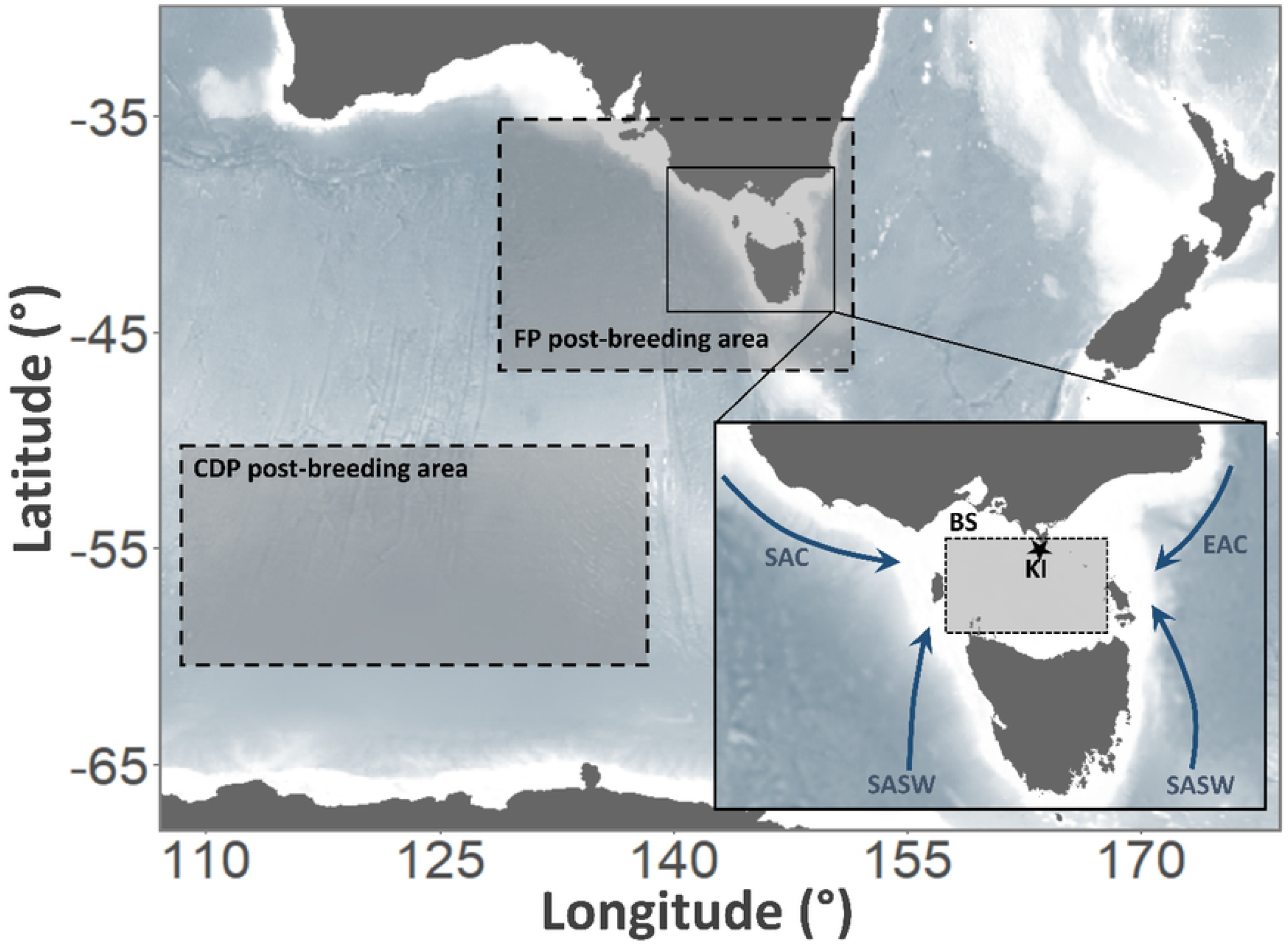
Study area of fairy prions (*Pachyptila turtur*) and common diving petrels (*Pelecanoides urinatrix*) from Kanowna Island, Bass Strait, south-eastern Australia. The lower right panel shows the main water masses influencing Bass Strait: Kanowna Island (KI); South Australian Current (SAC); Sub-Antarctic Surface Water (SASW); East Australian Current (EAC). The shaded rectangles describe the areas for which monthly sea surface temperature was extracted: Bass Strait (BS; study breeding region; Fromant et al. unpublished data); post-breeding area of fairy prion (FP Post-breeding area; Fromant et al. unpublished data); post-breeding area of common diving petrels (CDP post-breeding area; [37]).

Breeding phenology was designated into laying, hatching and fledging dates. Laying dates were considered as the first day the egg was found in the burrow if the burrow was checked regularly, otherwise laying dates were estimated from the observed hatching dates [46, 47]. The incubation period was denoted as the time from laying to hatching, the chick-rearing period was denoted as the time from when the chick hatched to when the chick was considered fledged [46, 47]. Fledging date was when a fully-feathered chick was no longer in the burrow.

Due to logistical constraints preventing access to the island at various times, the fledging dates of FP could not be recorded in 2017 and, therefore, these were estimated using previously-reported data on the duration of the chick-rearing period for the species [47]. Similarly, the laying and hatching dates for some CDP individuals could not be recorded in 2017 and were estimated using wing length-age relationships [45, 46, 48].

Once hatching was detected, to minimise disturbance to the nest, chicks were not handled during the brood stage (age 1-3 d and 1-10 d for FP and CDP, respectively; [46, 47]. Thereafter, chicks were monitored every 3-5 d when weather conditions allowed. To determine the growth rates of chicks, individuals were weighed in a cloth bag with a spring scale (± 2 g, Pesola Precision Scale, Schindellegi, Switzerland), bill and tarsus measurements were taken with Vernier callipers (± 0.1 mm) and wing length was measured with a shoulder-stopped ruler (± 1.0 mm). Mass at fledging was assumed to be the final measurement before the chick departed the burrow. Chicks no longer in the burrow that were too young to fledge were assumed to be dead if there were obvious signs of predation or if the burrow was invaded by a short-tailed shearwater (*Ardenna tenuirostris*).

Although most Procellariiform species do not show adverse effects related to repeated handling of chicks [49–51], the degree of response is likely to vary with environmental conditions and among species [51]. Therefore, the present study evaluated the potential impact of burrow disturbance and chick-handling in FP and CDP. During the 2018/19 breeding season, a group of control burrows (FP n = 20, and for CDP n = 16) was monitored monthly for each species and the hatching, fledging and breeding success were compared to those of study burrows.

To investigate the influence of environmental factors on breeding biology, monthly averages in sea surface temperature (SST) was obtained for areas considered important to the study species (Fig 1). The summer period plays a key role in the growth and reproduction of Australian krill (*Nyctiphanes australis*) in Bass Strait [52], the main prey of both FP and CDP [44]. Additionally, CDP have been observed to migrate during the post-breeding period to subantarctic regions [37] which may provide resources necessary for winter survival and preparation for subsequent breeding. However, FP stay in south-eastern Australia during the inter-breeding period and regularly visit Bass Strait islands during this period [43, 53]. Therefore, Bass Strait (38-40 °S, 144-148 °E; Dec-Nov), the post-breeding area of FP (35-48 °S, 130-152 °E; Feb-Oct) and the post-breeding area of CDP (50-60 °S, 110-140 °E; Dec-Apr) were used as proxies to investigate potential environmental influences on breeding in both species (Fig 1). All environmental data were downloaded from the Copernicus platform [http://marine.copernicus.eu/].

All statistical analyses were conducted in the R statistical environment 4.0.0 (R Core Team 2020). Data normality and homogeneity of variances were assessed with Shapiro-Wilk and Bartlett tests, respectively. Inter-annual variations in phenology (laying, hatching and fledging dates) were tested using analyses of variance (ANOVA or Welch’s ANOVA), and *post-hoc* tests were conducted using *t*-tests (parametric), or Kruskal-Wallis and Mann-Whitney *U* tests (non-parametric) depending on the data distributions. The growth curve analysis method [54] was used to investigate the potential variation in chick growth among years. Growth in body mass, wing length, tarsus length and bill length were modelled with a third-order orthogonal polynomial and fixed effects on all age terms. The model also included chick individual as a random effect. Hatching success (eggs hatched as a proportion of eggs laid), fledging success (chicks fledged as a proportion of eggs hatched) and breeding success (chicks fledged as a proportion of eggs laid) were compared between years using a Pearson’s chi-squared test.

## Results

For both FP and CDP, no significant differences were found for hatching, fledging or breeding success between control and experimental groups (Pearson’s Chi-squared test, all *P* > 0.4). Therefore, for both species, data from control and study groups were pooled to investigate differences between years. For the four study years, the FP laying period occurred between 10 and 31 October (Table 1). Mean laying date varied significantly between years (Kruskal-Wallis test: ***χ***^2^ = 57.456, *P* < 0.001), with eggs being laid significantly later in 2018/19 and 2019/20 than in 2017/18 and 2020/21. Correspondingly, mean hatching period was significantly delayed 2018/19 and 2019/20 compared to 2017/18 and 2020/21 (Kruskal-Wallis test: ***χ***^2^ = 51.939, *P* < 0.001), as well as fledging (Table 1; Kruskal-Wallis test: ***χ***^2^ = 47.28, *P* < 0.001). The average duration of the incubation and chick-rearing periods for FP were consistent between years at, 47 ± 1 d and 50 ± 1 d, respectively.

**Table 1:** Timing of laying, hatching and fledging dates (mean ± SE) of fairy prions (*Pachyptila turtur*) and common diving petrels (*Pelecanoides urinatrix*) from Kanowna Island, Bass Strait, south-eastern Australia. For each period/species, values not sharing the same superscript letter (a, b, c or d) are significantly different (Mann-Whitney *U* test: *P* < 0.05). Sample sizes are provided in parentheses.

| Species | Year | Laying | Hatching | Fledging |
| --- | --- | --- | --- | --- |
| Fairy prion | 2017/18 | 19 Oct ± 1 d (43)<sup>a</sup> | 05 Dec ± 1 d (43)<sup>a</sup> | 22 Jan ± 1 d (38)<sup>*</sup> |
| | 2018/19 | 24 Oct ± 1 d (51)<sup>b</sup> | 10 Dec ± 1 d (48)<sup>b</sup> | 27 Jan ± 1 d (29)<sup>a</sup> |
| | 2019/20 | 22 Oct ± 1 d (56)<sup>b</sup> | 07 Dec ± 1 d (43)<sup>ab</sup> | 24 Jan ± 1 d (28)<sup>b</sup> |
| | 2020/21 | 14 Oct ± 1 d (13)<sup>c</sup> | 30 Nov ± 1 d (14)<sup>c</sup> | 16 Jan ± 1 d (12)<sup>c</sup> |
| Common diving petrel | 2017/18 | 27 Jul ± 1 d (61)<sup>**</sup> | 20 Sep ± 1 d (61)<sup>a</sup> | 14 Nov ± 1 d (61)<sup>a</sup> |
| | 2018/19 | 15 Sep ± 2 d (43)<sup>a</sup> | 07 Nov ± 2 d (33)<sup>b</sup> | 25 Dec ± 4 d (9)<sup>b</sup> |
| | 2019/20 | 05 Sep ± 2 d (35)<sup>b</sup> | 02 Nov ± 4 d (25)<sup>c</sup> | # |
| | 2020/21 | 12 Aug ± 2 d (33)<sup>c</sup> | 06 Oct ± 2 d (33)<sup>d</sup> | 19 Nov ± 10 d (33)<sup>c</sup> |

\* dates estimated based on mean fairy prion chick-rearing period ( $50 \pm 1$ d)
\*\* duration dates estimated based on mean common diving petrel incubation duration

The breeding phenology in CDP was also significantly delayed in 2018/19 and 2019/20 compared to 2017/18 and 2020/21, though to a much greater degree than in FP (Fig 2). Mean laying date were 50 d in 2018/19 and 40 d in 2019/20 later than in 2017/18 (Table 1; Kruskal-Wallis test: ***χ***^2^ = 112.75, *P* < 0.001). Correspondingly, the hatching dates were 48 d and 43 d later in 2018/19 and 2019/20, respectively, than in 2017/18 (Kruskal-Wallis test: ***χ***^2^ = 89.26, *P* < 0.001), and fledging date was 41 d later in 2018/19 than in 2017/18 (*t*-test: *t_9.097_* = −11.092, *P* < 0.001; no study nests fledged in 2019/20). Intermediate phenology was observed in 2020/21, with the mean laying date being delayed 16 d compared to 2017/18 (Table 1). The duration of incubation was longer in 2018/19 and 2019/20 compared to 2020/21 (56 ± 1 d, 57 ± 1 d and 55 ± 1, respectively; Kruskal-Wallis test: ***χ***^2^ = 15.289, *P* < 0.001; no data available in 2017/18). Conversely, the duration of the chick-rearing period was not different between years (54 ± 1 d in 2017/18, 55 ± 1 d in 2018/19 and 55 ± 1 d in 2020/21, respectively; Kruskal-Wallis test: ***χ***^2^ = 2.081, *P* = 0.353; no study nests fledged in 2019/20).

**Figure 2:**
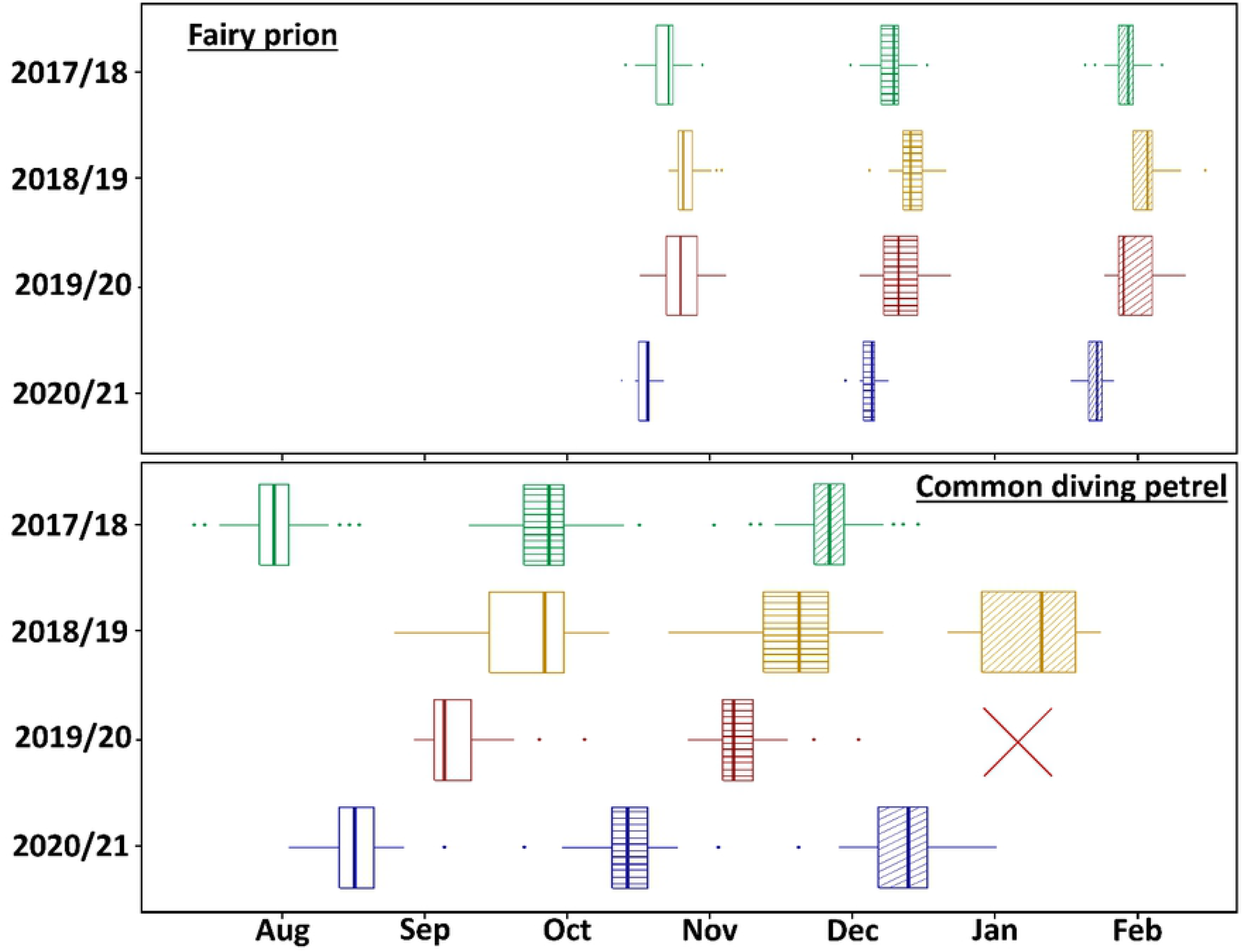
Inter-annual variation in the breeding phenology of fairy prions (*Pachyptila turtur*) and common diving petrels (*Pelecanoides urinatrix*) from Kanowna Island, Bass Strait, south-eastern Australia. Laying date = open boxplots; hatching date = horizontal dashed boxplots; fledging date = diagonal dashed boxplot. Laying date of CDP in 2017/18 was estimated based on mean incubation duration. Fledging date of FP in 2017/18 was estimated based on mean chick-rearing duration. Fledging date for CDP in 2019/20 was not determined because all study nests failed so no fledging data was available (red cross).

Hatching success for FP was not significantly different between years (Pearson’s Chi-squared test, ***χ***^2^ = 3.627, *P* = 0.305) (Table 2). Due to logistical constraints, no data were available for fledging and breeding success in 2017/18. However, the study nests were monitored during the early chick-rearing period (up to age 12 d) and the proportion of chicks surviving during this time was not significantly different between years (93% in 2017/18, 83% in 2018/19, 88% in 2019/20 and 91% in 2020/21; Pearson’s Chi-squared test, ***χ***^2^ = 1.795, *P* = 0.616). Fledging success (Pearson’s Chi-squared test, ***χ***^2^ = 4.688, *P* = 0.096) and breeding success (Pearson’s Chi-squared test, ***χ***^2^ = 3.713, *P* = 0.156) did not vary significantly between years.

**Table 2:**
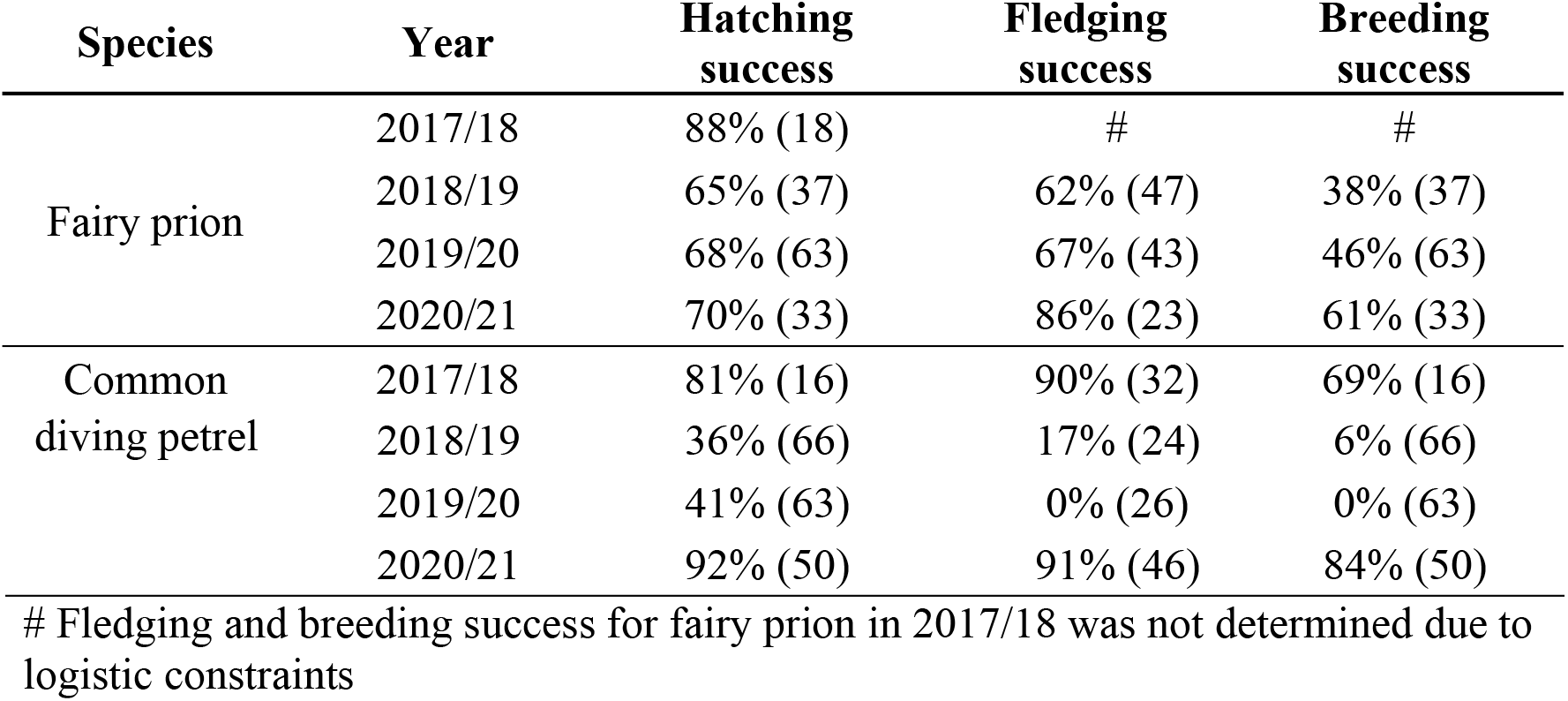
Hatching, fledging and breeding success (mean ± SE) of fairy prions (*Pachyptila turtur*) and common diving petrels (*Pelecanoides urinatrix*) from Kanowna Island, Bass Strait, south-eastern Australia. Sample sizes are provided in parentheses

Hatching success for CDP was significantly higher in 2017/18 and 2020/21 than in 2018/19 and 2019/20 (Pearson’s Chi-squared test, ***χ***^2^ = 46.291, *P* < 0.001) (Table 2). Similarly, fledging success varied significantly between years (Pearson’s Chi-squared test, ***χ***^2^ = 90.391, *P* < 0.001), fluctuating from 0% in 2019/20 to 91% in 2020/21. Correspondingly, breeding success in 2017/18 (69%) and 2020/21 (84%) was significantly higher than in 2018/19 (6%) and 2019/20 (0%) (Pearson’s Chi-squared test, ***χ***^2^ = 127.73, *P* < 0.001).

For both species, growth parameters (body mass, wing, tarsus and bill lengths) varied significantly among years (Fig 3, Table 3). There was a significant effect of year on both the intercept and quadratic terms, highlighting that growth rates decreased from 2017/18 to 2019/20. For FP, measurements of 12 d old chicks were significantly greater in 2017/18 and 2020/21 than in 2018/19 and 2019/20 (Fig 3). The difference was particularly pronounced between 2017/18 and 2019/20 for the wing length (*t*-test: *t_24.58_* = 5.803, *P* < 0.001) and tarsus length (*t*-test: *t_21.694_* = 2.337, *P* = 0.029). Additionally, body mass of FP fledging chicks was lower in 2019/20 than in 2018/19 (*t*-test: *t_29.333_* = 3.129, *P* = 0.004), similarly to wing length *t*-test: *t_35.965_* = 3.592, *P* < 0.001) and tarsus length (*t*-test: *t_32.699_* = 2.728, *P* = 0.010). For CDP, body mass at fledging was significantly higher in 2017/18 and 2020/21 than in 2018/19 (Table 3; *t*-test: *P* < 0.03), as was wing length between 2017/18 and 2018/19 (*t*-test: *t_4.269_* = 2.924, *P* = 0.039) but not in bill length (*t*-test: *P* > 0.161) or tarsus length (*t*-test: *P* > 0.323). In 2019/20, chick body mass growth rate decreased rapidly and lead to the death of all chicks during the first half of the chick-rearing period (mean age of chicks when found dead was 17.6 ± 8.2 d).

**Figure 3:**
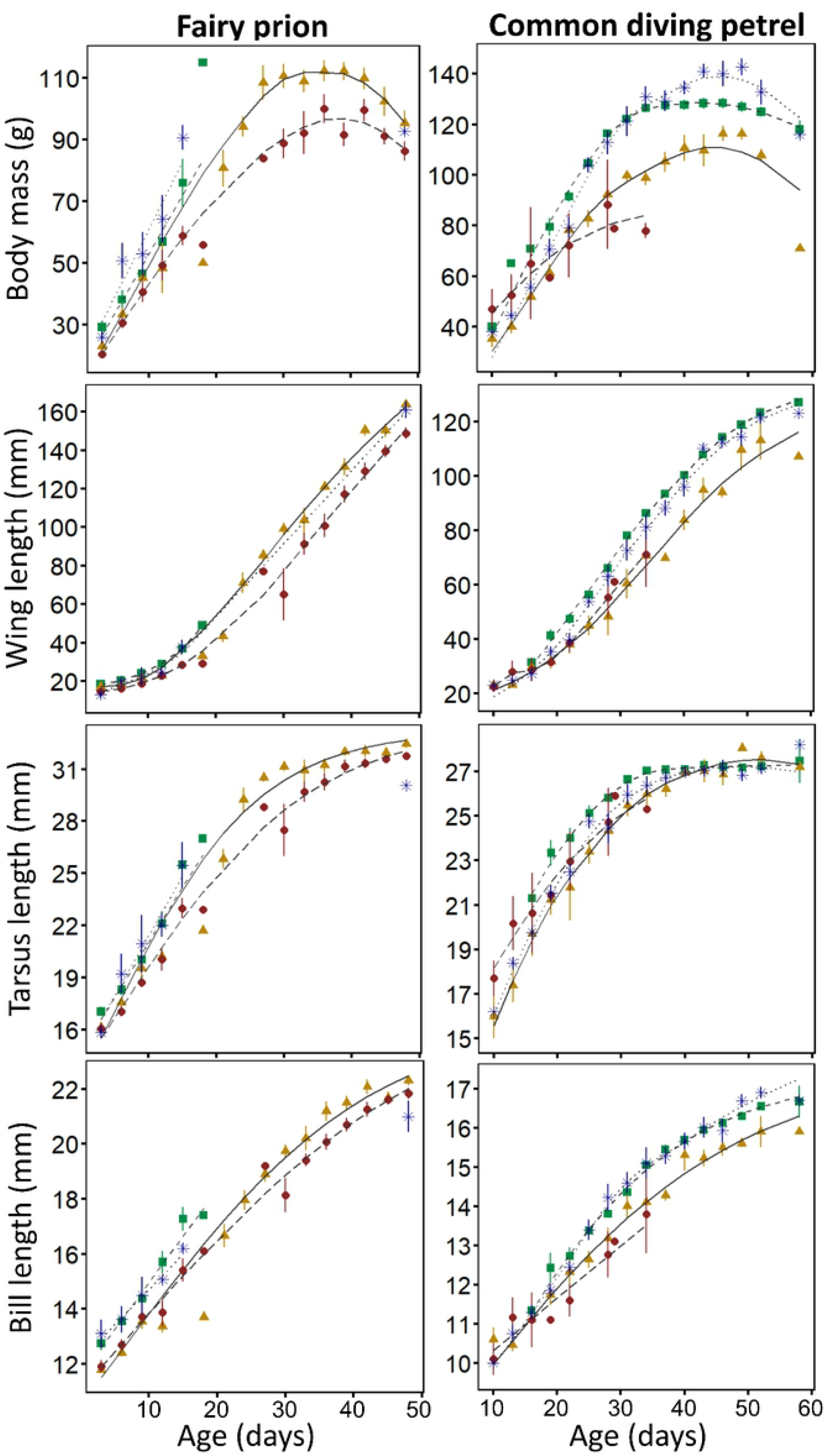
Growth curves of chick fairy prions (*Pachyptila turtur*) and common diving petrels (*Pelecanoides urinatrix*) from Kanowna Island in 2017/18 (green squares), 2018/19 (orange triangles), 2019/20 (red circles) and 2020/21 (blue stars).

**Table 3:** Body mass, wing length, tarsus length, and bill length (mean ± SE) of fairy prion (*Pachyptila turtur*) and common diving petrel (*Pelecanoides urinatrix*) fledglings from Kanowna Island, Bass Strait, south-eastern Australia. For each period/species, values not sharing the same superscript letter (a, b, or c) are significantly different (Mann-Whitney *U* test: *P* < 0.05). Sample sizes are provided in parentheses.

| Species | Year | Body mass (g) | Wing length (mm) | Tarsus length (mm) | Bill length (mm) |
| --- | --- | --- | --- | --- | --- |
| Fairy prion | 2017/18 (0) | # | # | # | # |
| | 2018/19 (23) | 99 ± 4<sup>a</sup> | 161 ± 2<sup>a</sup> | 32.4 ± 0.1<sup>a</sup> | 22.2 ± 0.1<sup>a</sup> |
| | 2019/20 (18) | 86 ± 2<sup>b</sup> | 142 ± 3<sup>b</sup> | 31.6 ± 0.2<sup>a</sup> | 21.8 ± 0.2<sup>a</sup> |
| | 2020/21 (2) | 92 ± 2 | 164 ± 6 | 30.1 ± 0.1 | 21.4 ± 0.7 |
| Common diving petrel | 2017/18 (40) | 126 ± 2<sup>a</sup> | 122 ± 1<sup>a</sup> | 27.3 ± 0.1<sup>a</sup> | 16.4 ± 0.1<sup>a</sup> |
| | 2018/19 (6) | 103 ± 8<sup>b</sup> | 113 ± 3<sup>b</sup> | 27.6 ± 0.3<sup>a</sup> | 16.2 ± 0.2<sup>a</sup> |
|  | 2019/20 (0) | ## | ## | ## | ## |
| | 2020/21(21) | 139 ± 5<sup>c</sup> | 118 ± 2<sup>b</sup> | 27.2 ± 0.2<sup>a</sup> | 16.6 ± 0.1<sup>a</sup> |

The SST in Bass Strait during summer preceding the breeding season of FP and CDP varied significantly between years (Dec-Feb; ANOVA: *F_10.976_* = 94.321, *P* < 0.001) and in the post-breeding migration area of CDP (Fig 4; ANOVA: *F_3.792_* = 18.803, *P* = 0.011). In Bass Strait, SST in the summers preceding the 2018/19 and 2019/20 breeding seasons was 2-4°C warmer than prior to the 2017/18 and 2020/21 breeding seasons. During this period, SST in Bass Strait was >1°C above the climatologic average for 95 consecutive days in 2018/19, and 68 consecutive days in 2019/20, while it was only four consecutive days above the average in 2017/18 and 0 in 2020/21, delineating 2018/19 and 2019/20 as years of marine heatwaves. Conversely, SST in the area used by CDP during the post-breeding migration exhibited an opposite trend, with January and February values prior to the 2017/18 breeding season higher than in 2018/19, 2019/20 and 2020/21 (Fig.4; ANOVA: *F_3.792_* = 18.803, *P* = 0.011).

**Figure 4:**
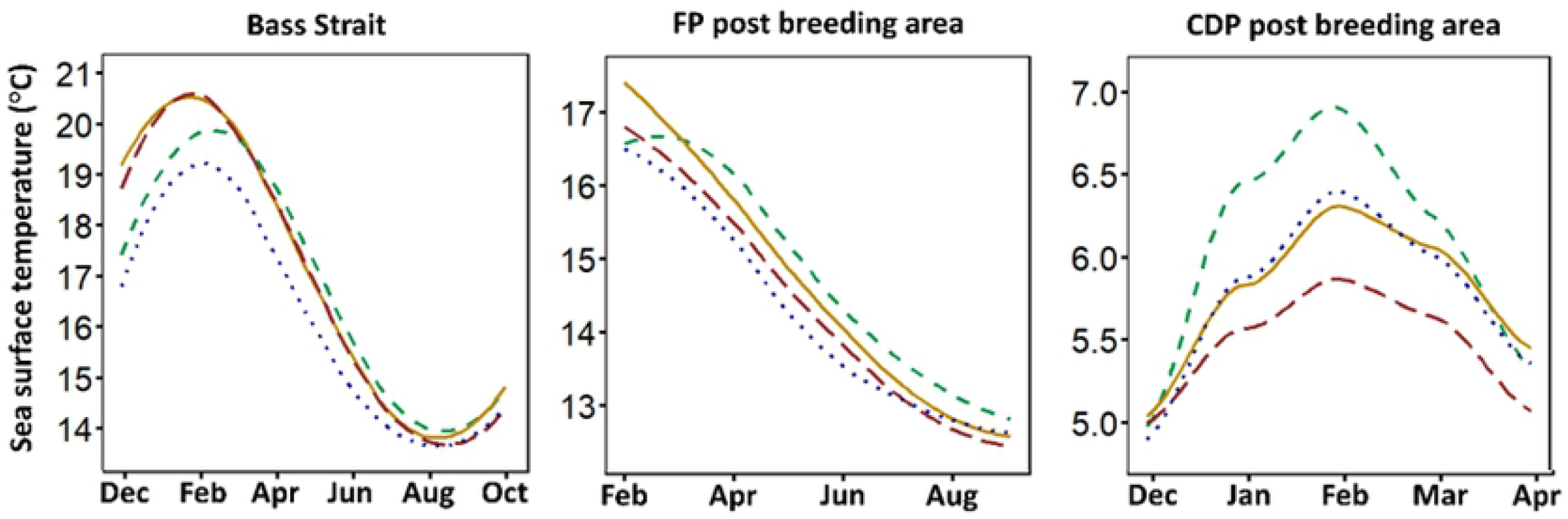
Inter-annual variation of sea surface temperature in three areas of importance to FP and CDP. The lines represent the conditions in the year preceding the 2017/18 breeding season (green dashed line); 2018/19 breeding season (orange line); 2019/20 breeding season (red dashed line); the 2020/21 breeding season (blue dotted line). Post-breeding migration area refers to the core area used by fairy prion and common diving petrels from Kanowna Island during their post-breeding migration (see Fig 1).

## Discussion

The present study documented inter-annual variations in the breeding phenology, chick growth and breeding success of FP and CDP breeding in northern Bass Strait, south-eastern Australia. Interestingly, these variations, which are likely to be related to shifts in food availability [55], differed between the species despite their consumption of similar prey [44]. The findings suggest potential differences in foraging areas and/or constraints on their breeding biology. Such observations from the northern extent of their range may provide insights into how these two ubiquitous species of the Southern Ocean may respond to anticipated effects of climate change in more southerly latitudes.

### Effect of environmental conditions on breeding

During the study period, both FP and CDP exhibited delayed phenology and low breeding performance in two consecutive years (2018/19 and 2019/20). Overlap between timing of reproduction and the peak of prey availability is crucial for seabird breeding success [56], with the timing of laying having been shown to be influenced by oceanographic conditions and, ultimately, food availability [57]. In the present study, the late start of the breeding season for both species in 2018/19 and 2019/20 was followed by substantial incubation failure, lower chick growth and/or lower fledging success and, for CDP in 2019/20, complete breeding failure. This suggests both species may have faced significant prey shortages during the two consecutive breeding seasons [58]. Additionally, for CDP, the rapid acceleration of breeding failure during the first few weeks post hatching suggests an energetic trade-off induced by a lack of food. With an exceptionally low prey availability, adults might not have been able to both self-maintain and feed their chick [59], leading to the extremely low breeding success [60]. As well as low food availability, low breeding success can be triggered by punctual extreme environmental events such as intense rain [61], or infectious diseases [62]. However, the sequence of delayed phenology, low hatching success, low chick growth and low fledging success support a scenario of reduced availability of important prey [63].

In south-eastern Australia, Australian krill plays a key role in the marine ecosystem with abundances observed to vary substantially between years, heavily affecting fish recruitment and seabird distribution and breeding success [64–66]. An optimal temperature for this large cold-water euphausiid ranges from 12-18°C [67] and a major factor influencing the survival and growth of Australian krill is ocean temperatures exceeding this range [64]. During recent decades, this optimal temperature has been disrupted in the region by persistently higher than average SST, otherwise identified as marine heatwaves [68, 69] which profoundly impact local zooplanktonic communities [70]. The marine heatwaves recorded in Bass Strait during the summers preceding the 2018/19 and 2019/20 breeding seasons, therefore, are likely to have affected the abundance of the main prey for both FP and CDP, inducing changes in their breeding biology. Additionally, the lowest SST recorded in the summer preceding the 2020/21 breeding season coincided with the highest recorded breeding success for both species and with CDP chicks reaching the highest maximum mass towards the end of fledging.

Variations in peripheral upwellings and currents around Bass Strait influence local oceanographic conditions, yet relatively little is known of the mechanisms involved. Warmer waters in Bass Strait can be the result of a lower influence of the northward cool Sub-Antarctic Surface Waters (SASW) and/or an increased eastward warm South Australian Current (SAC) and southward East Australian Current (EAC) [71]. Specifically, over recent decades, the strengthening of the EAC has resulted in a large influx of warm subtropical water southward [72, 73], inducing a replacement of large cool water zooplanktonic communities (including Australian krill) by smaller sub-tropical species [70]. Additionally, the strength and duration of the numerus upwelling systems on the edges of Bass Strait vary seasonally and annually [74], and their effect on the inner part of the strait depend on the confounding effects of the currents cited above [71].

Similarly, low food availability during the non-breeding period can induce low breeding performances through carry-over effects [75]. This period is critical for both study species, as adults need to restore their body reserves and, more importantly, undertake the energetically costly renewal of their plumage [37, 76]. In the post-breeding area of CDP in the Southern Ocean, zooplanktonic communities can be affected by warmer sea surface temperatures [77] impacting the body condition and survival of planktivorous seabird species [78]. As shown in Fig 4, the highest SST in the CDP post-breeding area corresponds to a year with a successful subsequent breeding season, while the lowest SST in the same area corresponds to the year of subsequent complete breeding failure. This suggests that the post-breeding area may not impose carry-over effects on the breeding season if local conditions are poor. In the present study, the link between zooplanktonic abundance and oceanographic variables remains unclear. Large-scale oceanographic indices such as the Southern Oscillation Index (SOI) and the Indian Ocean Dipole (IOD) have also been used to investigate the impact of temperature on the variation in planktonic productivity [79]. Observations in the present study show that marine heatwaves occurring in the years preceding each breeding season influences breeding biology, yet, the mechanisms causing these lags are unknown. Environmental variation may impact seabird species differently and further research exploring these mechanisms will provide knowledge about the responses of small seabird species that are in areas exposed to extreme oceanographic changes, like the Southern Ocean [25, 80].

### Different responses to environmental variability

In the present study, the impact of environmental variability differed greatly between the two study species. In contrast to FP, CDP exhibited inter-annual variations in breeding biology unexpectedly large for a subantarctic/temperate species [60]. The observed variation in phenology of 40-50 d far exceeds what has previously been reported in Procellariiformes and is highly unusual for temperate/subpolar species [81, 82]. Throughout their range, CDP have been shown to display variation in breeding phenology, with subantarctic populations laying in the mid-spring (October) [48, 83] while some populations have bred successfully during winter in Australia and New Zealand [46, 84]. Temporal shifts in breeding may affect the availability of food and nesting habitat as well as having implications for interspecific competition.

Additionally, there was no overlap between the chick rearing periods of both species during the years of high breeding success. In 2017/18 and 2020/21, the average CDP fledging date was 11-20 days before the average FP hatching date, reducing the possibility of interspecific competition for the same prey species in Bass Strait during the most energetically expensive period of breeding. In contrast, the chick-rearing period in both species overlapped by 15 days in 2018/19 and the mean death/failure date of CDP in 2019/20 was 18 days before the FP hatching date such that there was no overlap in chick-rearing. Although uncommon at the most northern extent of their range [46, 47], FP and CDP display full temporal overlap in breeding season throughout their subantarctic range [60, 85]. Avoiding an overlap in breeding season is a niche segregation mechanism for reducing interspecific competition in species that rely on similar prey. The temporal segregation in breeding season of CDP and FP observed in Bass Strait may enable both species to acquire local prey resources for chick-rearing without having to compete with each other. Therefore, when conditions lead to a delayed breeding season for CDP in Bass Strait, it could lead to increased difficulty for both species.

While FP and CDP feed mainly on the same prey during the breeding season [44, 60], their physiological and ecological differences may explain the contrasting impacts of environmental variability on these species. Provisioning chicks with highly concentrated stomach oil [86], FP chicks are able to regulate energy and water supply as well as allowing both adult and chick to endure long periods of fasting [86]. Being the only Procellariiformes to not produce stomach oil [34, 81], diving petrels chicks are subsequently required to be fed twice as frequently as other Procellariiformes [34]. Compared to CDP, the physiological characteristics of other small Procellariiformes are likely to provide better tools to cope with a decrease in prey availability in Bass Strait. In addition to the absence of stomach oil, the high wing load of CDP might be detrimental to their capacity to extend their foraging range and trip duration. Indeed, while CDP are restricted to short trips in both incubation and chick-rearing (< 200 km; 1-2 days; [18, 60, 87], FP are able to extend their range and trip duration to access distant foraging areas (600 km; 1-7 days) (Fromant et al. unpublished data).

Although FP appeared to respond better to environmental variations than CDP in the present study, they still experienced slower growth and poorer body condition of fledglings during breeding seasons following marine heatwaves. Such reduced fledging size and condition could lead to reduced first year survival [88]. Therefore, while the eco-physiological flexibility of FP may mitigate the apparent intensity and ecological impact of environmental variability [89], the long-term effects remain to be explored. Since CDP and FP share breeding periods during the austral summer throughout their sub-antarctic range, a delay in the breeding season further challenges breeding outcomes as winter snow and ice impact nesting and foraging options. Therefore, gathering long-term ecological and demographic knowledge about small Procellariiformes is particularly important to observe and understand the evolutionary response of Southern Ocean communities facing climate change.

## Acknowledgments

Fieldwork was conducted on Wamoon country, land of the Boon Wurrung, Bunurong and Gunaikurnai people. The authors thank the numerous field volunteers that took part in the data collection over the four field seasons (Fanny, Chloe, Louarn, Ruth, Lauren and Jess) and Sean for providing seamless access to fieldwork sites over the duration of the project.

